# Concentration invariant odor coding

**DOI:** 10.1101/125039

**Authors:** Christopher D. Wilson, Gabriela O. Serrano, Alexei A. Koulakov, Dmitry Rinberg

## Abstract

Humans can identify visual objects independently of view angle and lighting, words independently of volume and pitch, and smells independently of concentration. The computational principles underlying invariant object recognition remain mostly unknown. Here we propose that, in olfaction, a small and relatively stable set made of the earliest activated receptors forms a code for concentration invariant odor identity. One prediction of this “primacy coding” scheme is that decisions based on odor identity can be made solely using early odor-evoked neural activity. Using an optogenetic masking paradigm, we define the sensory integration time necessary for odor identification and demonstrate that animals can use information occurring <100 ms after inhalation onset to identify odors. Using multi-electrode array recordings of odor responses in the olfactory bulb, we find that concentration invariant units respond earliest and at latencies that are within this behaviorally-defined time window. We propose a computational model demonstrating how such a code can be read by neural circuits of the olfactory system.

---

A substantial computational challenge for the olfactory system lies in resolving odorant identities despite fluctuations in odor concentration arising from proximity to odorant source, air turbulence, and natural breathing^1-3^. Odorants are sensed by olfactory sensory neurons (OSNs), each expressing one out of a large family of olfactory receptor (OR) genes^4^. Axon terminals from OSNs expressing the same OR gene converge in a few discrete structures in the olfactory bulb (OB) called glomeruli. Odorants evoke responses in an ensemble of glomeruli to create a combinatorial representation of odor identity^5,6^. This representation varies not only across odorants, but also across concentrations of a single odorant^7-9^. Low concentrations of odorant evoke activity in only the most sensitive glomeruli, while increases in concentration result in recruitment additional less sensitive glomeruli. Despite this variability, odors’ qualitative identities are preserved across a range of concentrations^2,10,11^.

In several sensory systems, including olfaction, neurons have been shown to convert the strength of excitatory input into latency of response^12^ and it has been hypothesized that ORs with high affinity will depolarize OSNs earlier than those with low affinity^13-15^. This emerges as a result of multiple processes including intracellular signal integration^16^ and temporal dynamics of odorant concentration^17-19^. For air breathing animals, sniffing determines the temporal dynamics of odorant concentration in the nose, resulting in an affinity-defined sequence of OSN recruitment. While these sequences of recruitment vary across odorants, they have been shown to be mostly concentration invariant as changes in concentration preserve temporal rankings of ORs with different affinities and these latencies are proposed to encode information about odor identity^15,20,21^.

Here, we propose a strategy for concentration-invariant odor identification which uses a few earliest activated glomeruli in a coding scheme we call “Primacy Coding”. We assume that thefirst few glomeruli activated during a sniff are those receiving input from the most sensitive ORs for a given odorant. We propose that solely the members of this small set of early glomeruli encode odor identity. While this primary set varies between odorants, we assume that it is mostly preserved across concentrations of the same odorant. As concentration is increased, responses of less sensitive glomeruli are recruited later than the primary set, thereby maintaining the members of the early set and preserving encoded odor identity information (Fig 1B, C).

**Figure 1.**
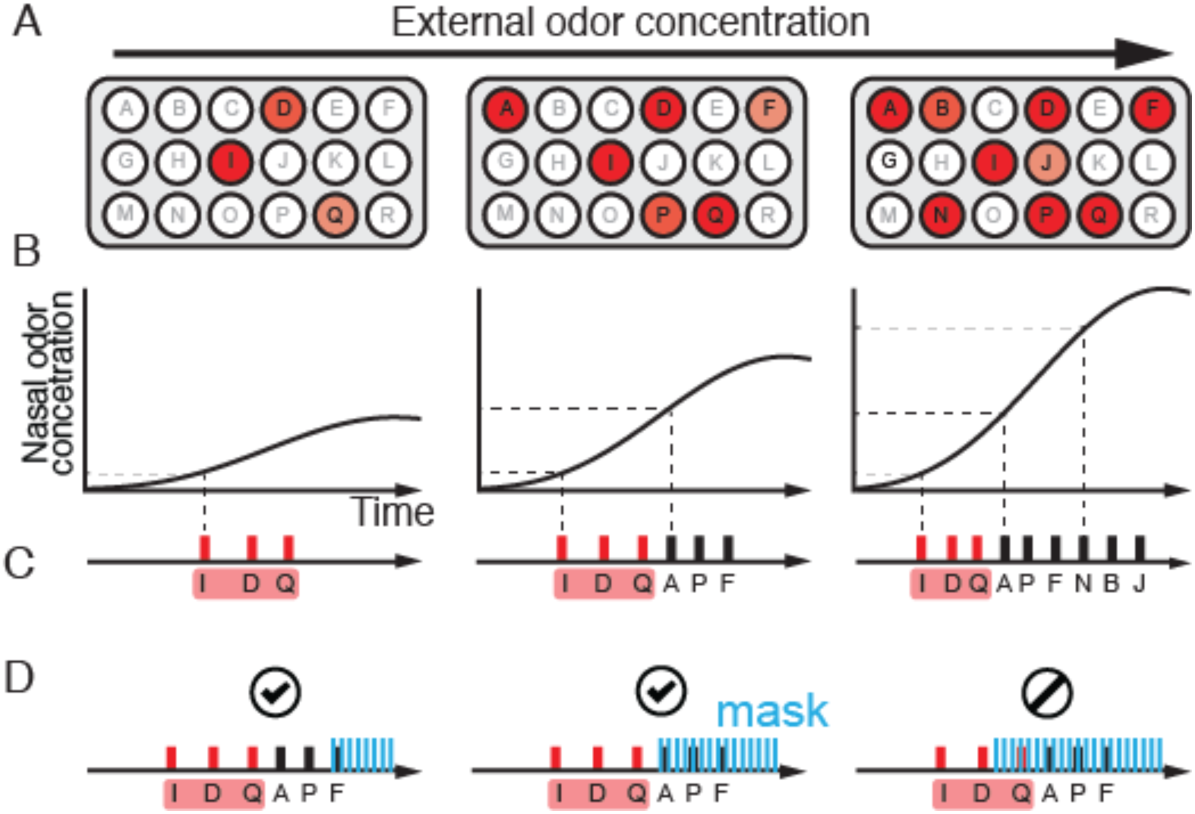
Primacy coding of odor identity. **A.** Schematic presentation of the patterns of glomerulus (OSN) activation for three concentrations of the same odor. The total number of active glomeruli increases with an increase of odor concentration. **B.** The temporal profiles of the odor concentration in the nose during inhalation, for three concentrations of the presented stimuli (dashed lines). **C.**Temporal sequence of glomeruli activation for three different odor concentrations. **D.** Optogenetic mask schematic demonstrating the effect of the optogenetic mask on temporal sequences when presented late and early relative to odor-evoked temporal pattern.

One of the central predictions of primacy coding is that animals use early “slices” of odor-evoked neural activity to define odor identity independently of the remainder of the pattern of evoked activity. To test this hypothesis, we developed an optogenetic masking paradigm in which we could create a temporally controlled masking stimulus during an odor discrimination task (Fig 1D). Through delayed triggering of this optogenetic masking stimulus relative to inhalation onset, we could preserve early epochs of odor-evoked information while making the overall combinatorial code unreliable through the activation of a large, heterogeneous subset of OSNs. We reasoned that if odor identity can be defined using only a small subset of early-responding primary glomeruli, our mask should not impair odor identification as long as it is initiated after this identity-defining subset. Conversely, activation of the mask before this initial subset of glomeruli become active should impair odor discrimination.

To produce the mask stimulus, we delivered 2 light pulses (25 mW, 2ms duration, 10 ms interpulse interval) to the olfactory epithelium in both nostrils of the transgenic mouse expressing Channel Rhodopsin2 (ChR2) in all OSNs^22^. The light was delivered via optical fiber stubs implanted above the olfactory epithelium. To characterize the neural response to the mask stimulus we measured light and odor evoked activity of mitral-tufted (MT) cells (n=119: 39 single-unit, 80 multi-unit) in the OB, which are the first recipient of information from OSNs (Fig.2A). The masking stimulus generated response occurring after a short delay following light onset (mean=11.5ms, mode=8ms) (Fig. 2C). The overall mask excitatory response lasts approximately 50 ms, followed by prolonged inhibitory response until approximately 200 ms (Fig. 2B, Extended Data Fig. 2E).

**Figure 2.**
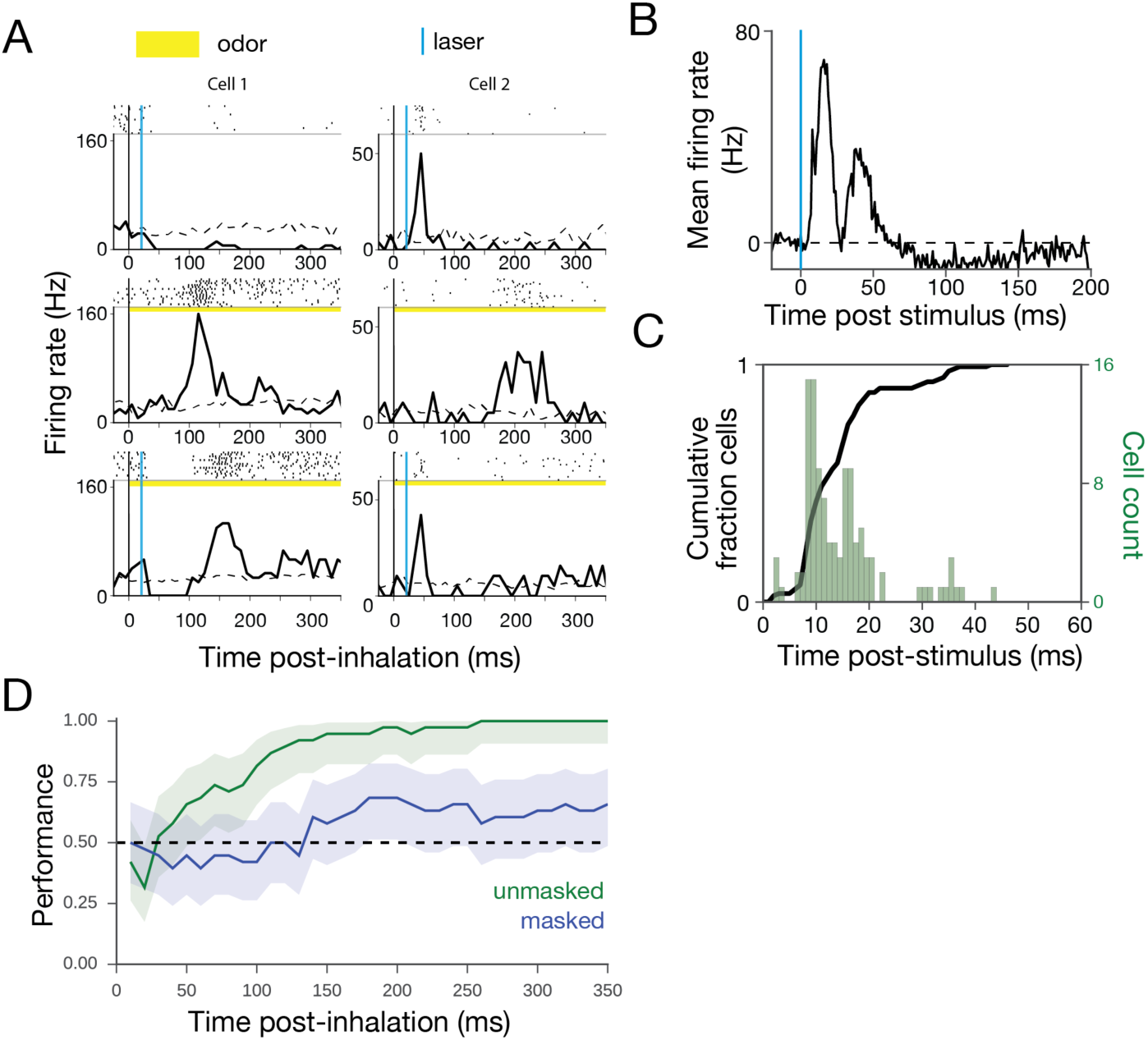
Neural response to optogenetic mask and effect on odor information. **A.** Example raster and PSTH for two example MT cells. Inhalation onset corresponds to t=0. Top: Example MT cell responses to optogenetic mask (25 mW, ON:2 ms − OFF:8 ms − ON:2ms), light stimulation at 20 ms post-inhalation. Middle: Odor response (left: 2-hydroxyacetophenone, right: alpha-pinene). Bottom: Odor plus laser mask. **B.** Baseline-subtracted mean of PSTHs for laser-responsive units (n=113). **C.** Cumulative distribution function (black line) and histogram of units’ response latencies to masking light stimulus. **D.** Linear classifier cross-validation results for unmasked data for two odor presentations (limonene, pinene). Responses to unmasked odor presentations were time-binned (10 ms bins) and used to train support vector machine (SVM) classifiers. For each time point, SVMs were trained on the response vector inclusive of bins from t=0 to that time. Each classifier’s performance is described by cross-validation on unmasked trials (blue) and testing on masked trials (green). Shaded areas indicate 95% confidence interval (Clopper-Pearson method).

How effective is the mask in eliminating odorant responses? Out of 119 recorded units, 29 responded to one of two odorants (pinene, limonene). The mask, presented at 20 ms latency post inhalation onset, modified most of the odor responses, even though odor responses typically occur much later than the mask stimulus (Fig. 2A). To characterize the effect of mask on information available for discrimination, a support vector machine (SVM) was trained to classify MT cell responses to two odorants (limonene vs pinene) with and without masking stimulus.

Based only on the odor responses of 29 units, classification performance on an individual trial raised from chance level to 100 *%* in the first 260 ms from the inhalation onset. On the masked trials, performance of the classifier stayed at chance level until ∼180 ms post inhalation onset, and never exceeded 68% (Fig. 2D). We may assume that the mask is efficient in eliminating odor information for at least 100 ms duration following the mask.

To test the effect of the mask on behavior, mice (n=3) were trained in a head-fixed 2-alternative choice paradigm to discriminate between two odorants (eugenol and 2-hydroxyacetophenone) for a water reward (Fig. 3). To ensure that decisions were based on odor identity and not intensity, we scrambled the odorant concentrations by presenting five concentrations within a two order of magnitude range. On probe trials within the session, the optogenetic mask was presented with target odor. The probe trial structure was used to prevent animals from adopting a novel strategy to overcome the effect of the mask. The animals’ performance was strongly affected by the masking stimulus when it was initiated between 0 and 50 ms after inhalation onset (Fig 3B, S1E). The presence of the mask lowered the mean performance at these early latencies to almost chance level of 56% compared with the unmasked performance of 92% (at odorant concentration 1 μΜ). As the onset latency of the mask is increased between 50 ms, performance in the odor identification task recovers and approximates the unmasked asymptotic performance at approximately 100 ms.

**Figure 3.**
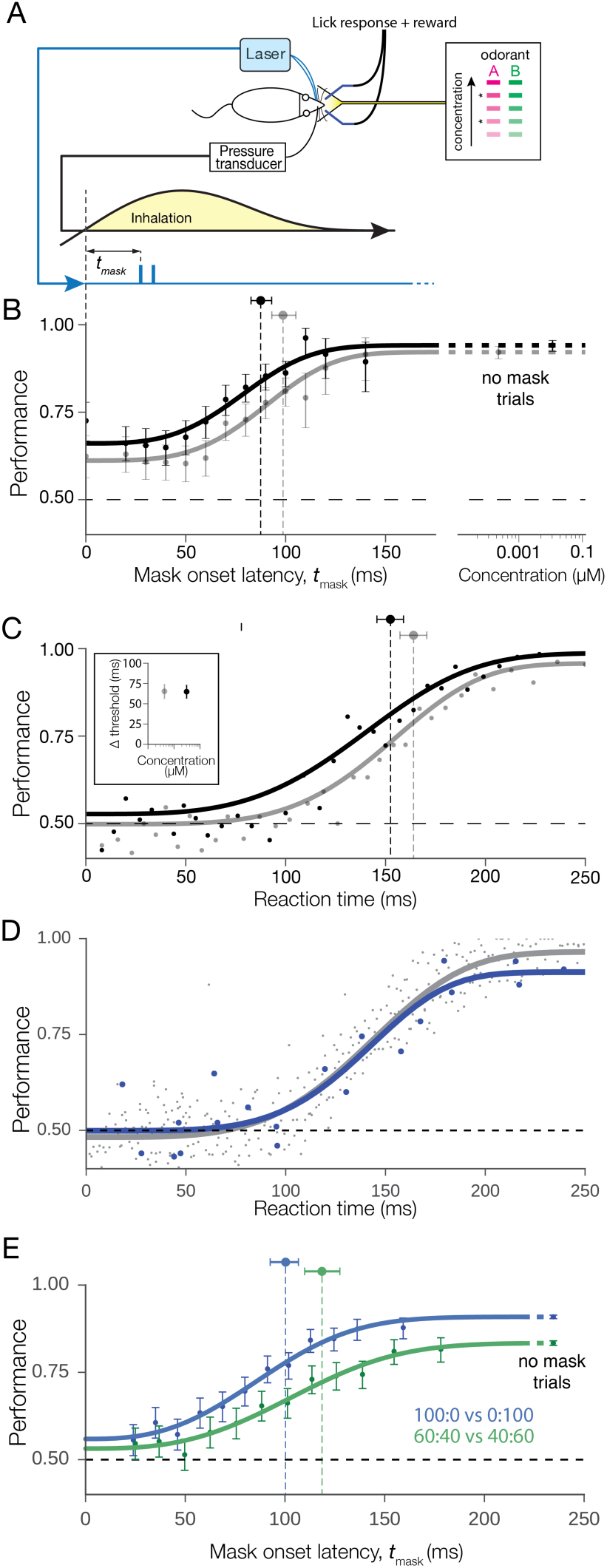
Optogenetic masking behavioral paradigm. **A.** Behavioral task schematic: Mice were trained to respond to 2 odors A and B at 5 different concentrations to lick left or right water spout. Mask timed to the onset of the first inhalation after odor delivery was presented during subset of trials for two concentrations (asterisks). **B.** Discrimination performance versus mask latency for odors (2-hydroxyacetophenone and eugenol) at two concentrations. Mask stimulus presentation was initiated on the first inhalation of odorant and after the mask onset latency, *t*_*mask*_, had elapsed. High concentration annotated in black markers, low concentration in grey. Error bars indicate 95% confidence interval estimates. Weibull fit to data indicated with thick lines. Markers above performance curves indicate Weibull threshold latency values for each fit with 95% confidence interval estimates. **C.** Mouse reaction time vs. performance in unmasked stimuli of two concentrations (as above). Dots indicate data binned into bins of 125 trials. Lines indicate Weibull fit. Inset: difference between mask and reaction time threshold latencies. **D.** Mouse reaction time vs. performance for unmasked trials compared with late masked trials (t >= 100 ms) for the same dataset. Points represent bins of 50 trials each. **E.** Performance versus mask latency for discrimination of pure (blue) carvone enantiomers versus mixtures made with those odorants (green).

Changes in odorant concentration should change the absolute timing of OSN recruitment and, thus, affect the timing of odor percept formation in our task. We tested this prediction by fitting a Weibull generalized linear model (see Methods) to masking data for two concentrations of odorant and comparing the thresholds of these fits. A 10-fold decrease in concentration delays the recovery of performance in masking trials by 13.3 ms (100.3 vs. 87.0 ms, bootstrapped 95% confidence intervals [94.6, 106.3] and [82.5, 92.1], one-sided bootstrap test p = 0.0007). As expected, odor percepts are formed later for lower concentration odorants, likely due to delayed recruitment of OSN activity^12^. Very similar dependencies have been observed for other odor pairs and 5 other mice (Fig. S1D).

Reaction time (RT) has been historically used to determine the timing of sensory processing and decision-making based on sensory stimuli. The dependence of performance on both mask latency and RT are qualitatively similar, except that the reaction-time curve is shifted by approximately 66 ms. This relationship is similar for both concentrations tested here (65.5 and 66.2 ms, Fig. 3C, Insert). The concentration-dependent shift in RT vs performance can be wholly explained by the shift observed in our masking paradigm, indicating that peripheral encoding of odor information limits the timing of olfactory decision making. The observed delay between the RT and mask can be ascribed to a motor delay that is constant across concentrations.

Is it possible that the presence of mask trials makes an animal to adopt decision strategy amplifying significance of the initial time interval? To address this issue, we performed two analyses. First, we compared reaction time in unmasked trials and trials where the mask was presented late (t ≥ 100 ms) and had minimal effect on performance (Fig. 3D and S1G). The Weibull fits to the dependencies of performance on reaction time in both cases are nearly identical. Second, we see no effects of learning on mask trials between early and late behavioral sessions, comparing the performance on the first and last 20 masked trials across all sessions (Fisher-Exact test p > 0.05). (Fig. S1F). Together, these results provide evidence that randomly presented masking trials do not change the animal decision strategy and animals typically use early evoked information in the odor discrimination tasks.

Does primacy coding strategy apply only to simple tasks? To make the task more difficult, we trained animals to discriminate between mixtures of carvone enantiomers (60:40 vs 40:60). Using mixtures to reduce the contrast between stimuli decreases discrimination performance relative to pure carvones (83.3% vs 90.8%) and delays the mask threshold time (118 ms vs 101 ms) (Fig. 3E). As with the latency shift between concentrations of odorant, the shift due todecreased contrast is consistent with the measure of reaction time vs performance (Fig. S1E). To generate an equivalent level of performance in the more difficult mixture discrimination task, sensory information must be integrated by the subject over a longer period of time. However, the extension in integration time is relatively small on the scale of the length of a sniff, demonstrating that animals are still using early information to make this more difficult discrimination.

Our behavioral results define the temporal window in which odor information is integrated. To determine if primacy codes exist in the OB at timescales consistent with our behavior, we recorded responses of 338 MT cells to 3 odorant concentrations spanning a range of 2 order of magnitude. The primacy coding model predicts the existence of MT neurons that display excitatory responses to odor across a wide range of concentrations in a narrow temporal window at the beginning of the sniff. Among 119 units which exhibit excitatory responses to odorant α-pinene, 15 responded to all three concentrations, a subset that we called “concentration-stable” (Fig. 4A). The remaining 104 units responded only to subset of concentrations, which we called “concentration-unstable” (Fig. 4B). As expected, a number of responsive units grew as concentration increased (Fig 4C). While the population response at low concentration was dominated by concentration-stable units, unstable units were 3.7 times more numerous than stable at the highest concentration tested.

**Figure 4.**
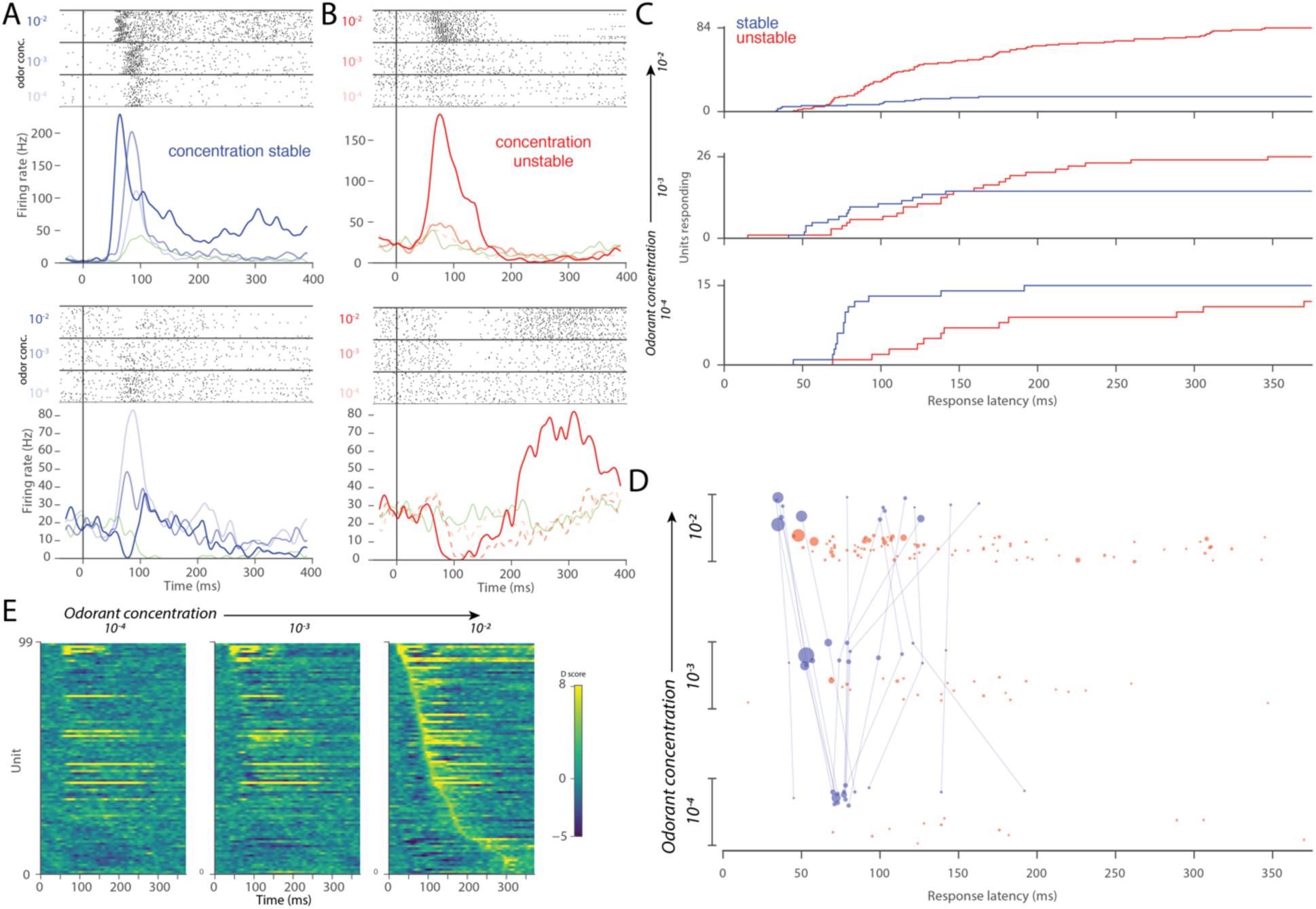
Concentration invariant MT cell response latencies correspond with behavior. **A.** Example raster and PSTH of two MT cells which were responsive to odorant across 3 orders of magnitude concentration. T=0 corresponds to inhalation onset. **B.** Same plots for MT cells which respond to only a subset of concentrations. Dashed PSTH lines represent responses that did not cross the threshold for significance (see methods). **C.** Cumulative distributions of MT cell response latencies. **D.** Response latencies of MT cell-odor pairs across concentrations for concentration stable and unstable units. Units’ responses across concentrations are connected. Area of each point in the plot is proportional to the Cohen’s D-score of the response over non-odor response (see methods). Point positions are randomly jittered on the concentration axis for visibility. **E.** Odor responses sorted by units’ response latencies at the highest concentration of odorant. Responses are represented as D-scores, where positive and negative scores are excitation and inhibition relative to blank inhalation responses.

According to our primacy coding model, the identity of early, stable units that are responsive across concentrations represent odor identity. The stable population’s mean response latency was shorter than the unstable population at all concentrations (conc 10^−3^: 86.8ms vs 177.2ms, p=0.0003; conc 10^−2^: 82.7ms vs 148.2ms, p=0.0071; and conc 10^−1^: 85.7ms vs 149.0ms, p=0.0249; one-sided KS test). While the latencies of the unstable and stable populations overlapped, a subset of stable units responded earlier than all unstable units at the highest concentration, consistent with the primacy coding model (Fig 4D). The latencies of these stable units scaled with concentration, as expected by the model and behavioral result. The mean latency shift between concentrations for this early, concentration-stable subset was 15.5 ms, (σ: 5.5 ms). This is comparable to the timing shifts in behavioral identification across concentration (13.3 ms). The stable population’s activity encoded odor-identity information and was not representative of non-specific odor responses; only one of these concentration-stable units was responsive to another odorant tested, α-limonene.

Importantly, we find that latencies and even latency relationships are not preserved across concentrations in awake animals, although this has been observed in anesthetized animals (Fig. 4D)^20^. Instead, both absolute and relative latencies of stable responses units shift dramatically as concentration increases, clustering into early and late subsets at the highest concentrations. We observed that the excitatory responses of late set of stable cells was proceeded by transient inhibition at high concentrations (Fig. 4A lower panel, 4E). This transient odor-evoked inhibition was observed widely in our recorded population (Fig. S3F, S3H), and we hypothesize that it is due to lateral and feedback inhibition in the OB. For late stable units, we predict that earlier members of the MT population drive inhibition arriving earlier than or in coincidence with feedforward excitation. This shunts the neuron’s initial excitatory response, causing an apparent increase in response latency. As a result, we conclude that these units are not in the primary set, as it is obvious that earlier members of the population must proceed them to generate the network activity responsible for this inhibition. So, while short latency is predictive of concentration-invariance, the relationships of response latencies within our recorded population as a whole are not conserved across concentrations and are unlikely to encode odor identity.

What elements of olfactory networks are sufficient to process primacy information? How does mask affect odor recognition? Our behavioral data shows an incomplete suppression of animal performance at small masking delays (Fig. 3B). In a more difficult discrimination task, the effect of mask is slightly delayed compared to easier tasks (Fig. 3E). Are these features expected within the primacy coding mechanism? To address these questions, we developed a computational primacy decoding model based on known features of the OB and piriform cortical (PC) circuits. First, our model includes random feedforward connectivity between the OB and PC, which provides the basis for coincidence detection of MT cell activity arriving early in the sniff cycle. Second, our model includes random recurrent inhibitory circuits in the PC (blue in Fig. 5A). The role of this connectivity is to suppress late arriving, “non-primary” input to PC neurons. This architecture is consistent with observed global and broadly tuned inhibition in the PC^23^ and has been used as a feature of network models for the processing of fine temporal information in the PC, including the proposed mechanism for coincidence detection^24^. Finally, we assumed that PC neurons have a memory property such that once activated, they can maintain the persistent activity in order to retain the odorant identity in the network until the initiation of action. Although our computational model does not explicitly specify the mechanism of persistency, it has been hypothesized to emerge from voltage dependent synaptic channels, such as NMDAR or GABA_B_ activating KIR channels^24^. Overall, our network included randomly connected 300 MC and 1000 PC neurons.

**Figure 5.**
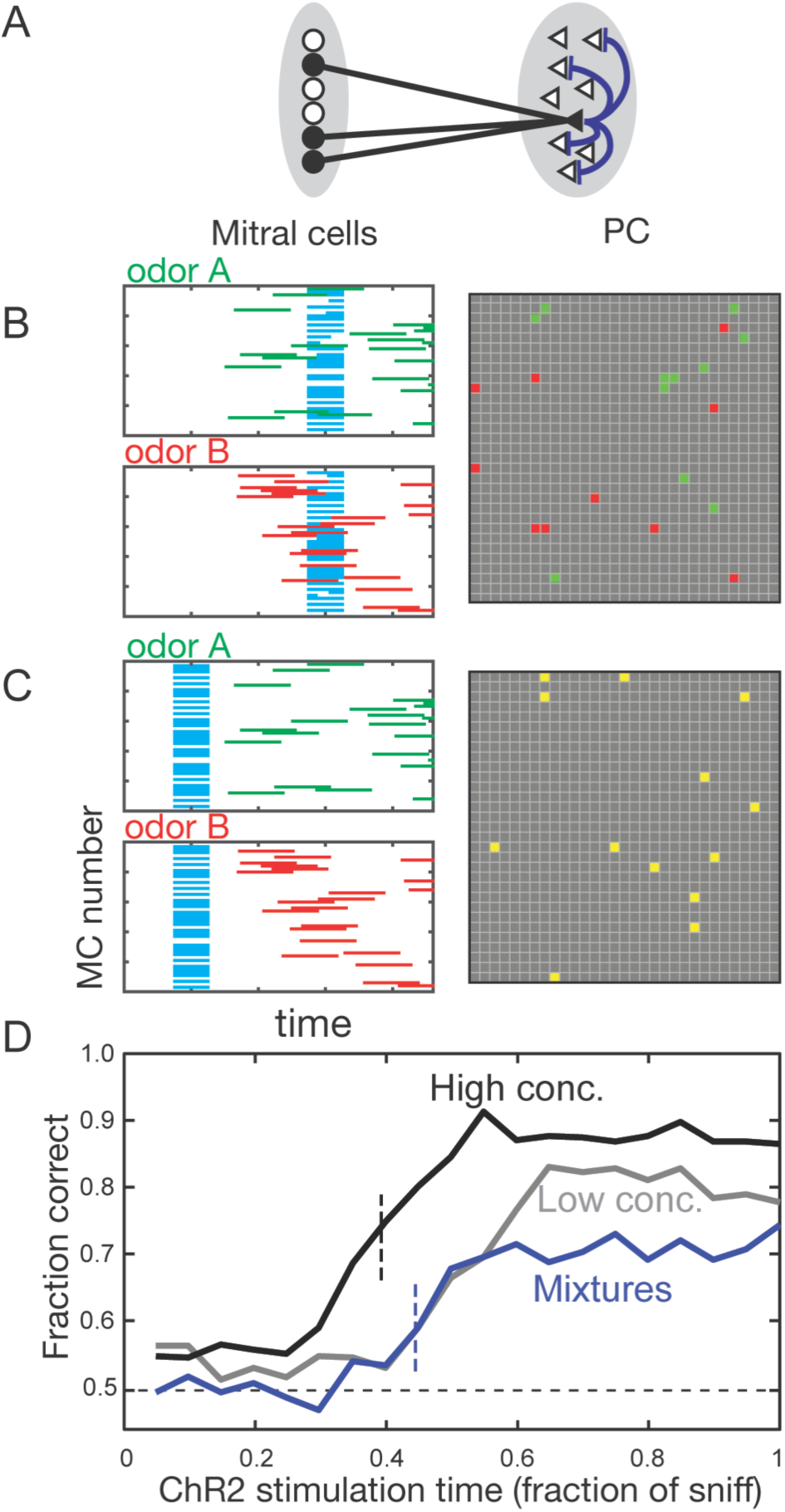
Computational model of primacy decoder can explain behavioral effects of mask. **A.**The schematic of the network included in the computational model. Pyramidal neurons in PC receive random excitatory connections from MT cells (black lines) and connected to a random subset of other pyramidal neurons via inhibitory connections (blue lines). The inhibitory connections are responsible for blocking of the inputs to PC after the initial (primary) representation is formed. **B-C.** Responses of MCs (100 out of 300) and PC cells for early (B) and late (C) masks. MCs respond to Odor A, Odor B, and ChR2 (green, red, cyan). PC responses at the end of sniff cycle (gray squares) are shown for two odors in the same panel. For the early mask (C), PC responses are indistinguishable for two smells (yellow). **D.** Discrimination success rate as a function of mask latency for three conditions: odors A vs. B – high concentrations (black line), odors A vs. B – low concentrations (gray line), odor mixtures 60%A+40%B vs 40%A+60%B (blue line). Low/high concentration conditions were introduced by expanding/compressing the timing of MC odorant-related inputs within the sniff cycle. Mixtures were introduced by combining together high conc. sequences of inputs corresponding to odors A and B. The temporal dependencies reproduce the basic qualitative features of data (Fig. 2B&E), including concentration-dependence of the error rate at different ChR2 timings.

Our model provides insights into the mechanism of optogenetic suppression of the animal’s performance for early delivered masks. We simulated odor-evoked activity in MT cells as a random spatiotemporal pattern and the mask as synchronous activity independent of odor (Fig. 5B,C). When the mask follows initial odor-evoked activity, it does not affect odorant dependent activity patterns in PC (Fig. 5B), due to the broad inhibitory network, which suppresses excitatory inputs from the late mask or odorant dependent inputs. In contrast, when the mask precedes odor activation, a pattern of activity emerging in the PC is not odorant-dependent and is different from those of the odorants alone (Fig. 5C). This light-evoked pattern can be viewed as a new percept that is unrelated to the original odors.

Our model qualitatively explains an incomplete suppression of discrimination for early masks (t<0.2) (Fig. 2B and Fig. 5D, black and gray lines). This occurs due to the presence of noise: on certain trials, odor-dependent OB responses are more robust than on average, causing failure of the mask on these trials (Fig. S4). In case of complex mixture, the behavioral response is delayed compared to an easy stimulus (Fig. 5D, blue vs. black lines) consistent with experimental results (Fig. 2E). This occurs because the differences in pools of primary glomeruli emerge slightly later in the sniff cycle, as both of the stimuli have similar sets of initially activated glomeruli. This result implies that primacy read out mechanism does not take the sequence of glomerular activation into account, since such differences are expected to occur early. In a version of the model sensitive to the activation sequence, the delay in performance for complex tasks is strongly reduced (Fig. S5). Overall, our computational model confirms that the experimental data is consistent with the identity of primary MT cells rather their recruitment order being important for the coding of odor identity.

We demonstrate here that the earliest evoked neural activity can be used to make olfactory decisions and demonstrate that neural activity in this time window may encode odor identity across concentrations through primacy coding. Previous behavioral studies demonstrate that early epochs of odor-evoked activity are sufficient for the detection of odorants^25,26^ and concentration discrimination^27^ in rodents. However, because these experiments did not scramble odor concentrations during testing, it is impossible to rule out whether animals were solely using intensity cues to guide their decisions. In earlier imaging studies it has been proposed that the most sensitive glomeruli may encode individual odors^15,28^, however, to our knowledge, our work provides the first behavioral evidence and computational support for this hypothesis.

While primacy coding relies on timing of activation of glomeruli, it has distinct features and predictions compared to latency coding schemes previously proposed in olfaction and other sensory modalities^14,24,29-31^. In fact, we find in our recordings that both absolute and relative latencies within the population of odor-responsive MT units are not maintained across concentrations in awake animals. Rather, primacy coding suggests the unique role of the early activated glomeruli, which we find animals can use and which appear to be stable across concentrations of odorant.

Second, the primacy model emphasizes the role of the set of early responding neurons rather than the sequence of glomerular activation. Our experiments with mixtures in combination with computational modeling provide indirect evidence towards this prediction. Further experiments with precise temporal control of early activated glomeruli may confirm this prediction.

Third, the primacy model sets limits for information capacity of the olfactory code. The upper bound estimate for the number of odorant identities represented by the primacy code is ∼*N*^*p*^/*p*! where N is the number of different OR types and p is size of the subset used to define identity. With p = 5−6 and the N = 350 OR genes found in the human genome, the coding scheme can represent ∼10^10^−10^12^ different odors.

Forth, the primacy model emphasizes the role of individual glomeruli for odor coding. In mice, a deletion of a single receptor TAAR4 is sufficient to abolish aversive behavior to a specific odor^32^. Studies of human genetic variability lends evidence that only highly sensitive receptors predominate in defining odor quality. Subjects with different alleles of a single OR report differences in perceptual qualities for strong ligands of the OR, while they are likely to report similar qualities for weaker ligands^33^.

Fifth, primacy makes specific predictions about mixtures of odorants because the earliest activity should dominate perceptual qualities of the mixture. Perceptual masking of slowly perceived odors by fast odors (temporal suppression) has been observed in human psychophysics, but warrants more attention^34^. If both odorants evoke early activity within the primary set, we predict that this combination will cooperate to synthesize a new odor percept. Finally, primacy coding does not make explicit provision for parallel processing of components in mixtures, a task where humans demonstrate poor performance^35^.

Primacy coding suggests a relatively simple solution to the complex computational problem of robust concentration invariant representation of odorant identity in olfaction. It provides inherently rapid odor identification, a vast coding capacity and can be implemented by the architecture of the olfactory system.

## Materials and Methods

### Mice

Behavioral concentration series data were collected in 4 OMP-ChR2-YFP heterozygous mice (2 female, 2 male). Mixture data were collected in a separate cohort of 5 OMP-ChR2-YFP heterozygous mice (2 female, 3 male). Electrophysiological data to characterize masking were collected from a separate cohort of mice (n=2). Five male C57B/6 mice (Jackson Labs) were used for concentration series electrophysiology. Subjects were 8-12 weeks old at implantation and were maintained on 12hr light-dark cycle in isolated cages after implantation. All procedures were approved by the IACUC of NYULMC in compliance with the NIH guidelines for the care and use of laboratory animals.

### Sniff recording

Sniff was monitored via intra-nasal pressure. An 8 mm long, 21-gauge cannula was implanted into the anterior dorsal recess. Total insertion depth from the surface of the nasal bone was 1.5 mm. Pressure change relative to atmospheric pressure was measured using a pressure sensor (24PCEFJ6G, Honeywell) and amplified (AD620, Analog Devices). A Schmitt (dual-threshold) trigger was used to define inhalation and exhalation onsets in real time on an Arduino microcontroller. For concentration series electrophysiology, respiration was measured using an externally mounted pressure sensor placed in front of the subjects’ nares.

### Surgery

Mice were anesthetized using isofluorane gas during surgery. The head bar, pressure cannula and optical fibers were implanted in a single surgery. The nasal cannula was implanted in a small hole in the anterior nasal bone and affixed with glue and dental cement. The optical fibers were implanted bilaterally in two holes drilled in the posterior nasal bones and affixed using the same technique. Two small screws (size #000-120 × 0.0625”, Small Parts, Inc.) used to stabilize the implant and provide electrical connection for lick detection were implanted in the skull at a location approximately corresponding with S1 cortex. Animals were allowed to recover for at least 3 days before water deprivation.

### Stimulus delivery and behavioral control

Behavioral control and data acquisition was computer-controlled using the custom Voyeur software^36^.

For odor stimulus delivery, we used an 8-odor olfactometer (Extended Data Fig. 1B). Odorants were diluted in mineral oil and stored in amber volatile organic analysis vials. Olfactometery manifolds, valves, and tubing contacting odorized air consisted entirely of PTFE to minimize cross-contamination of odorant. During odor presentation, nitrogen carrier gas is diverted through a single vial and enters the main air stream, resulting in an adjustable dilution in a range between 10x and 100x. Airflow rates for carrier and main flow rates were controlled using two mass flow controllers (Bronkhorst). During periods between stimuli, animals were presented with 1 L/min background air and olfactometer air was directed to exhaust using a four-way PTFE final valve (NResearch).

Odorant concentrations were controlled using a combination of gas- and liquid-phase dilution. Through manipulation of the ratio of odorized and clean air flowrates, we were able to achieve a dilution range of 10-1% odorized air (10x). In experiments where more that 10-fold dilution was required, we used liquid dilution to increase our range. Because liquid dilution ratios do not accurately predict headspace (gas-phase) concentrations, liquid dilutions were assayed using a photo ionization detector (Aurora Scientific) to determine relative concentrations of odorant in gas-phase. For concentration-series electrophysiology, all dilutions were made in air-phase by diluting odorized air with unodorized air.

Mixtures of carvone enantiomers were made in liquid phase. Enantiomers have identical vapor pressures and solvent interactions, thereby allowing accurate prediction of component ratios in gas phase from liquid mixture ratios.

Light masking stimulus was provided by two 473-nm, 105 um ID fiber coupled diode laser (Blue Sky Research, PN: FTEC2471) terminated in a ceramic ferrule. During behavioral sessions, the laser source ferrule was mated to a ferrule permanently implanted on the mouse. Implanted ferrules (MM-FER2007-304-4050, Precision Fiber Products, Milpitas, CA) were fabricated with 400 um ID, 0.39 NA fiber (FT400UMT, Thorlabs, Newton, NJ) and etched using hydrofluoric acid to provide diffuse light within the nasal cavity. Laser power was calibrated using a light power meter (Thorlabs) prior to behavioral sessions at the ferrule tip.

Water reward stimulus was delivered through two 21-gauge stainless steel lick tubes (Small Parts) and controlled using pinch valves (BioChem Fluidics).

Licks were detected by measuring change in resistance at the lick port when animals made contact with the lick tube using custom lick detectors (Janelia, HHMI).

### Behavioral procedure and training

Animals were water deprived for at least 5 days prior to start of behavioral training. Animals were housed on a 12:12 light-dark cycle were tested between ZT 15 and ZT 24. To acclimatize animal to head-fixation and behavioral apparatus, animals were shaped by given water through a single lick tube until animals received their entire 1 ml water ration during a session. In subsequent sessions, a second lick tube was introduced. To encourage exploratory behavior in subsequent training, animals were rewarded for alternating licks between left and right lick tubes. Two-lick shaping sessions persisted until animals successfully received entire 1 ml water ration in a session.

Odor discrimination was trained on subsequent sessions with only slight modifications to the paradigm used in testing sessions. After a variable inter-trial interval (12-15 seconds) and with the condition that 1 second had elapsed since the last licking activity was recorded. Odor stimulus was delivered for 500 ms and was initiated on the start of exhalation so that odor stimulus was stable prior to inhalation (Extended Data Fig. 1C-D). For each trial, odor concentration was drawn randomly. After stimulus onset, a “grace” period was enforced where licks were not scored to reduce impulsive licking prior to odor sampling. To eliminate stereotypic response bias, trials were chosen using a bias correction algorithm during training and testing^37^. For initial training, this grace period included time point until 500 ms after the first inhalation of odorant. After criterion performance was met, this grace period was shorted in the following session to 300, then to 150 ms for testing. Randomization was preformed using Mersenne Twister RNG (Numpy).

Testing sessions were conducted only after animals reached criterion performance (>80%) on the odor discrimination task. Mask trials was randomly interleaved in sessions for only 2 of the 5 concentrations presented (Fig. 2A). Trial ordering within a session was computer controlled and the investigator was blind to the conditions of each trial. For masking data at multiple concentrations, masking trials was presented in every other session with both concentrations masked in the same sessions at rate of 17% of total trials. For masking data for carvone mixtures, masking trials were interleaved throughout each session at a rate of 8.3% of total trials. Data were excluded from animals that did not complete training and testing due to loss of sniff signal, illness, or loss of implant. Two animals were excluded from mixture experiment due to faulty fiber implantation. Data from experiments represented in Fig 3B and S2D were collected from the same animals.

### Behavioral analysis

All behavioral analysis was conducted using custom scripts on the Anaconda Python distribution (NumPy 1.9.2, SciPy 0.15.1)^38^. Binomial proportion confidence intervals were calculated using the Clopper-Pearson “exact” method. Trials from sessions in which animals preformed a level of <80% correct responses were excluded from analysis. Data were fit with the Weibull psychometric function using maximum likelihood estimation method:

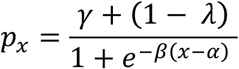

For masking data,γ (guess rate: the asymptotic performance at short latencies) was fixed using the average from masking at time points < 60 ms and λ (lapse rate: the asymptotic performance at long latencies) was fixed based on the data obtained in unmasked trials within these sessions. For reaction time analysis, all parameters were fit. Confidence intervals for fit parameters (thresholds) were estimated using the 2.5^th^ and 97.5^th^ percentile of distributions created by fitting each of 10,000 bootstrap simulations for each experimental condition. To bootstrap, trials were randomly drawn with replacement using Mersenne Twister RNG (Numpy).

Reaction time data was taken only from trials in which no mask stimuli were presented. These sessions were interleaved with mask sessions. Reaction time performance data and timing was taken using the first responses following odor stimulus onset irrespective of grace period. These data were fit using the same techniques as above, but with all parameters free. Trials with very long reaction times (>= 300 ms) were truncated from this analysis, as performance was not monotonic after approximately 300 ms.

### Masking electrophysiology

Electrophysiology was conducted in awake animals during using acute recording techniques. Six-shank, 64-ch silicone probes (Buzsaki 64sp, NeuroNexus) were used to record neural activity. Neural data were acquired using “Whisper” acquisition system (Janelia, HHMI) at 20833 Hz using SpikeGL software (Janelia, HHMI). Action potentials were detected and clustered using SpikeDetekt2 and KlustaKwik with manual clustering preformed using KlustaViewa^39^.

All basic analysis was done using the Anaconda Python distribution. Mask response latency was determined by comparing baseline (no mask) activity distribution to mask response. To construct baseline sample distribution, PSTHs for 7 sniffs prior to every mask trial were sampled. From these baseline samples, 100,000 samples of the same size as the number of mask trial were drawn with replacement to create a simulated PSTH from the same number of trials as the masked PSTH. Finally, the PSTH from masked trials was compared with the baseline PSTH distribution. Latency to mask response was defined as the first bin where firing rate was >3 fold greater than the baseline PSTH and the bin was at the 0.0001th percentile of the bootstrapped baseline distribution.

### Linear SVM classifier

Population vectors were assembled from spike trains of recorded unit (n=29) that responded to one of the odors presented. For each cell and for each trial a 35-dimensional vector was created by binning action potential events into 10 ms bins from the 0 ms to 350 ms after the first inhalation onset of the odorant. Activity vectors from cells were concatenated and standardized to make a population vector for training and testing. Linear classifiers were created using the Scikit-learn v0.16.1 LinearSVC class^40^. To assess performance of the classifiers on unmasked trials, a leave-one-out cross validation strategy was used. To classify masked trials, classifiers were built using all available unmasked trials (N=19), and the classifier was scored on its performance at classifying all masked trials.

### Concentration series electrophysiology

NeuroNexus A64 Poly5 2x32 probes were used to record acutely from awake animals. Units were detected using Spyking Circus software (v0.3.0)^41^. Significant odor responses were defined by comparing inhalation-aligned blank odor response distributions with responses to odor presentations (Fig. S3A-D). To establish these distributions, blank and odor responses were bootstrapped across trials, summed across trials, and smoothed with a 3-sigma Gaussian kernel of width 30 ms. For each time point, the bootstrapped firing rates were fitted with a Gaussian and the Cohen’s D (discriminability) score was calculated for each time bin. Response latencies were defined as the first time-bin in with a D-score above 5, where overlap is < 0.01 (Fig. S3E).

### Computational model

Our model was based on random and sparse connectivity between the MT cells and the PC cells, as well as within the cortex as detailed below. Our simulation included 300 MT cells and 1000 cells in the PC. Neurons in PC formed random sparse inhibitory recurrent connections to other cells in PC with 50% probability. Non-zero inhibitory recurrent connection weights within PC were 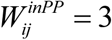. MT cells also formed random excitatory connections with the PC cells with 13% probability (each PC cell received inputs from 40 MTCs). Non-zero projections from MTCs to a PC cell were 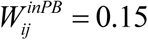. In the case of order dependent model, we used 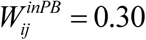.

The state of each PC neuron was defined by the input that this neuron receives *ui* that satisfied the equation 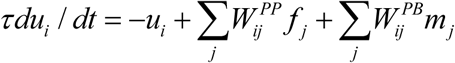. Here τ = 0.05 is the time constant measured in the fraction of sniffing cycle. The activation state for each PC neuron had a hysteretic dependence on its inputs *f*_*i*_ = *F*_±_(*u*_*i*_). The activation function *F*_±_ was single valued for values of input variable *u* satisfying *u* > *u*_+_ = 0.2 and *u* < *u*_−_ = −150. For these values of parameters, the activation function *F*_±_ was equal to 1 and 0, respectively. Within the bistable range, i.e., for *u*_−_ ≤ *u* ≤ *u*_+_, *F*_±_ was bistable and remained constant depending on prior history. Therefore, if a neuron was activated, the activation function within the bistable range remained equal to 1, whereas for an inactivated neuron, the activation function was 0. Activation occurred when inputs exceeded *u*_+_, and inactivation happened when inputs fell below *u*_−_.

Each simulation was carried out over the period of 700 time units using Runge-Kutta method with time step Δ*t* = 0.002. The simulation started at t = −0.2 and lasted until t=1.2. t=0 corresponds to the onset of inhalation, and t=1 is the end of the early stage of the sniff cycle. The masking stimulus was presented between t=0 and t=1. This time interval was expected to reproduce the early part of the sniff cycle during which odorant identity is established.

To model MT cell responses to odorants we generated random spatiotemporal patterns of MT cell activity. For the case of pure odorants (black and gray in Fig. 5D), we used two different random patterns for each of the two odorants. The response of each responsive MT cell was represented by a transient that lasted 0.5 time units (in fractions of the early part of the sniff cycle). The transient response consisted of an increase of the output of a mitral cell *m*_*j*_ from 0 to 1 and reset back to 0. For low/high concentration conditions, the earliest transients began at t=0.4/0.25 respectively and the following MTC transients were distributed at random times with the time step of 0.02 (every 0.02 a MTC was recruited). This feature was intended to replicate the tendency of MT cells to respond later in the sniff cycle to lower odorant concentrations^42^. For the mixture case (Fig. 5D), we generated two sets of random recruitment orders for each of the pure binary compounds within the mixture and combined them into a single sequence by offsetting one of them by 2 positions, depending on which component concentration was bigger (60% + 40% vs. 40% + 60% composition). Following the earliest transient onset at t=0.25, as in high concentration case, one MTC was recruited every time interval of 0.02 with the recruitment order as described above. In each trial, MTC transients had a finite probability to be emitted to mimic the experimentally observed transient event reliability^43^. The probability was p=0.8/0.9 for low/high concentration conditions. To simulate the ChR2 stimulation, we added a pulse that began at the time indicated in Figure 4 and lasted 0.1. The amplitude of the pulse was 0.18. The pulse was present in 75% of MTCs chosen randomly. We added normally distributed white noise with the standard deviation of 0.1 to the activity of MTCs. This was done to reproduce observed features of psychophysical performance. We tested our simulations for a range of parameters and verified that the qualitative conclusions are not sensitive to the exact set of parameters chosen. We ran 500 trials for each of the two odorants and each concentration. After each set of 10 trials, we reset the randomly selected weights and parameters in the model to mimic trials performed by different animals.

The perceived identity of the stimulus in each trial was inferred from the activity of PC cells at the end of the simulation (t=1.2) from the template pattern of PC response that maximally overlapped the evoked response. To obtain the template, we ran the simulation once without noise for every condition.

#### Code availability

Code used for data acquisition is available at https://github.com/olfa-lab. Code used for analysis and modeling will be made available upon request.

## End Notes

### Acknowledgements

We would like to acknowledge Thomas Bozza, Matt Wachowiak, Kevin Franks and Andreas Schaefer for commenting on this manuscript; Admir Resulaj for his technical assistance in engineering the behavioral apparatus; and Ezequiel Arneodo and Kristina Penikis for critical discussions and for sharing the design of apparatus used for electrophysiological recordings. This work was supported by NIDCD (R01DC013797, R01DC014366) and the Whitehall Foundation. C.D.W. is funded by N.S.F GFRP.

### Author Contributions

C.D.W. and D.R. designed behavioral and electrophysiological experiments; C.D.W. and G.O.S. collected behavioral data; C.D.W. collected electrophysiological data; C.D.W. analyzed experimental data; A.A.K. created computational model; D.R., C.D.W., and A.A.K. wrote the manuscript. All authors discussed results and commented on the manuscript.

### Author information

The authors declare no competing financial interests. Correspondence and request for materials should be addressed to D.R.

## Supplementary Figures

**Supplementary Figure 1.**
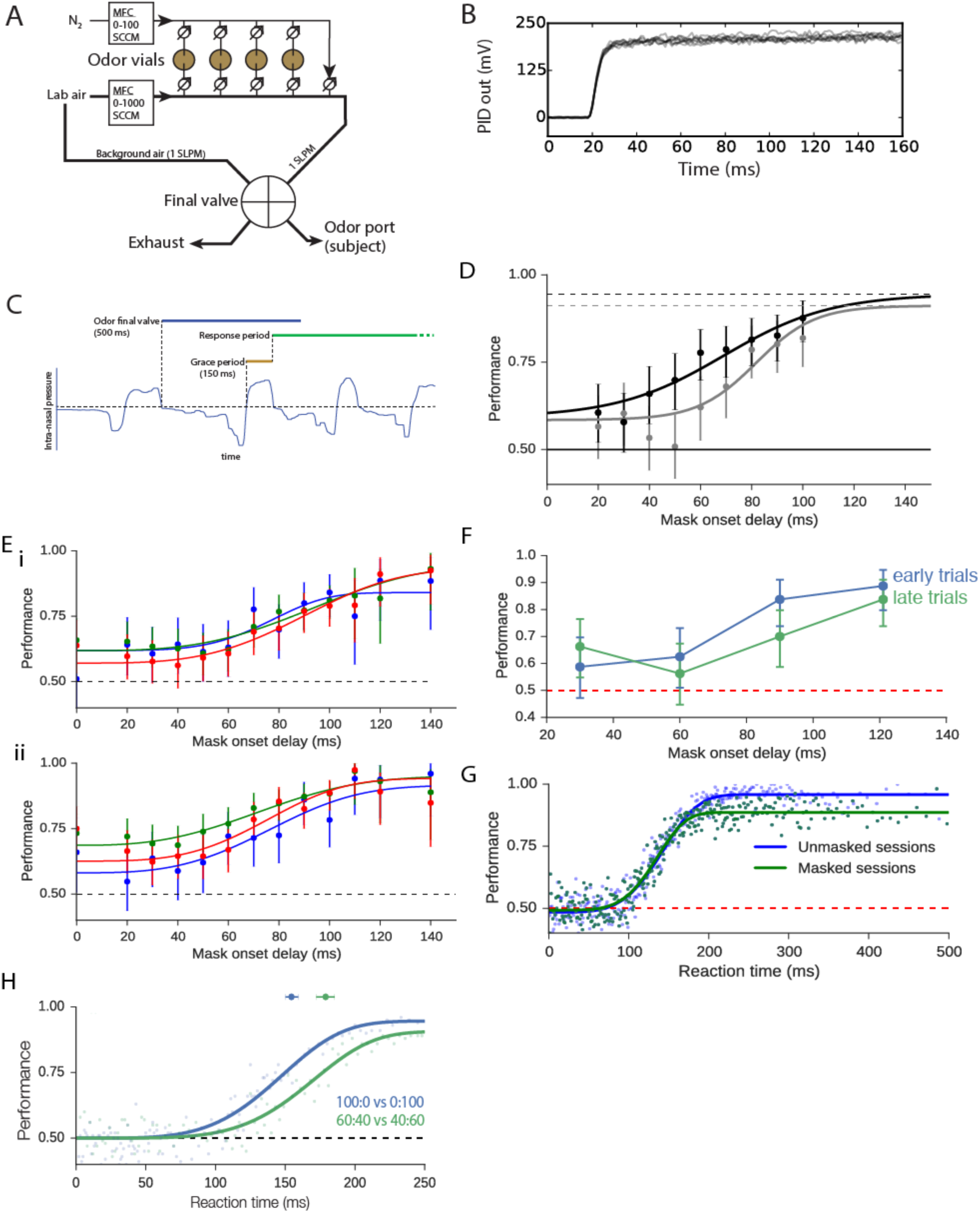
Behavioral apparatus and experiments. **A.** Olfactometer schematic. Illustration of 4-vial olfactometer. Liquid-diluted odorant is carried by nitrogen gas (N_2_) flow into parallel carrier air stream when two vial valves are opened. Only one vial is exposed to nitrogen/air streams at a time to prevent cross-contamination. Sum of N_2_ and air streams is 1 SLPM controlled by 2 mass flow controllers (MFCs), and adjusting ratio of these gasses controls dilution of odorant. Final valve (FV) positioned downstream of olfactometer is used to ensure fast and stable onset of odorant and continuous flow 1 LPM air for subjects. **B.** Photoionization detector trace of odor time course demonstrates fast and repeatable odor presentations. FV is triggered on at t=0. D) Subjects to first inhalation following FV onset. Less than 0.5% of trials (*n* = 66,765) had inhalation before odor was available to the subject (30 ms). **C.** Schematic of 2AFC behavioral trial. Behavioral trials are initiated with odor final valve opening at the start of the first exhalation following inter-trial-interval. Lick responses from are recorded but ignored until 150 ms after the initiation of first inhalation after odor valve trigger (grace period). Response period is relatively unrestricted (total trial time 2.5 seconds). **D.** Performance vs mask onset delay for additional odor set (limonene vs pinene) at two concentrations: 0.1% (grey) and 1% dilution (black). Mask parameters were (100x1ms pulse, 200 Hz, 25 mW) for these experiments. Data were pooled for 4 subjects. Error bars express 95% confidence intervals. **E.** Data and Weibull fits for individual subjects from Figure 3B for low (i) and high concentrations (ii). **F.**Comparison of performance on early and late masking trials. The first 20 and last 20 masked trials for each animal at different delay time points were analyzed. Because of a low number of trials, data were pooled for all subjects (n=4) and data were binned for time points within 30 milliseconds. X axis value denotes the high value of the bin for each data point. **G.**Comparison of reaction time to performance for sessions with and without masked trials. **H.** Performance versus reaction time for pure carvone discrimination and carvone mixture discrimination.

**Supplementary Figure 2.**
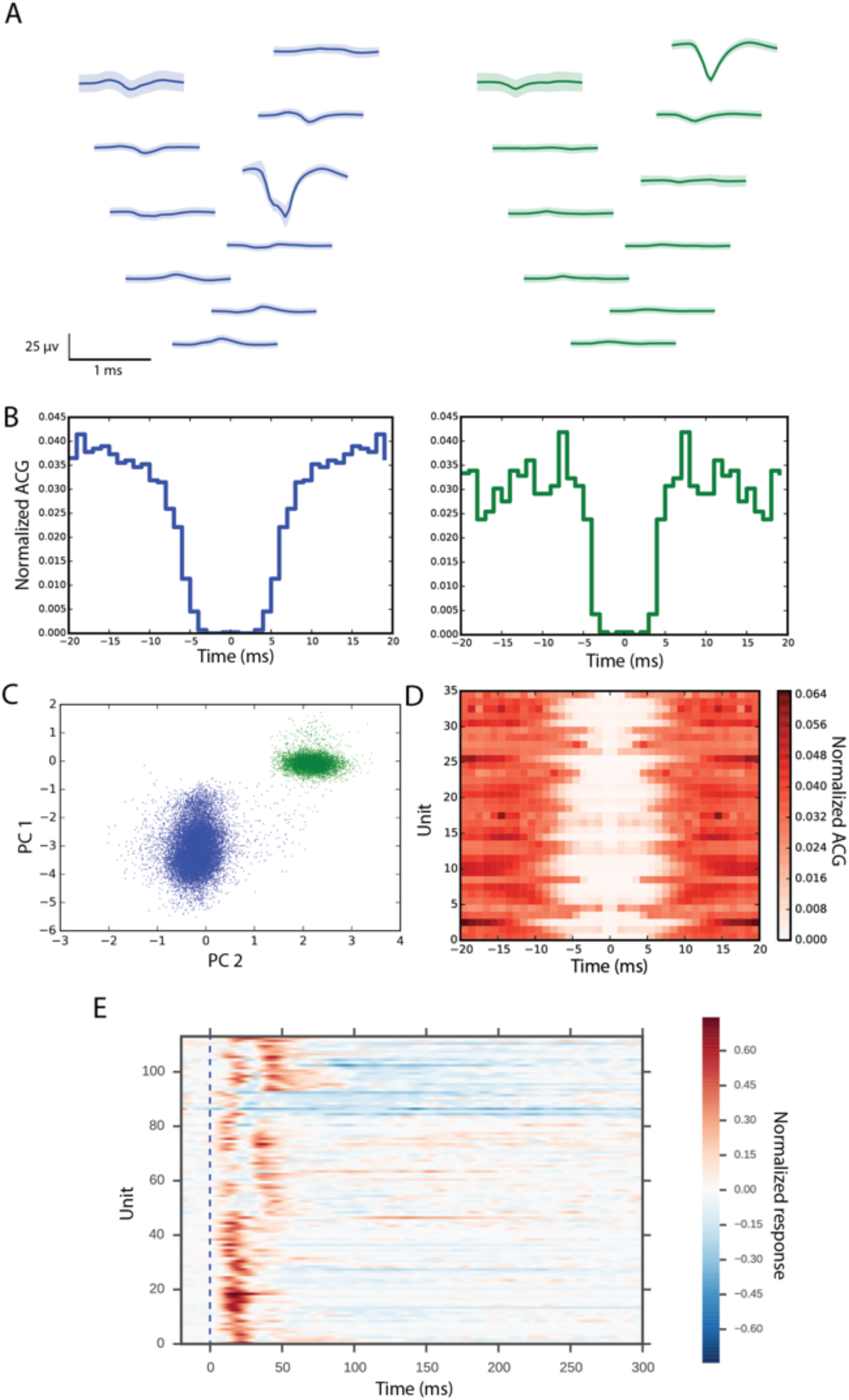
Masking electrophysiology supplementary information. **A.** Average waveforms of two single units (SU) recorded units recorded on the same electrode shank. Waveforms are displayed for all of the electrode sites on the shank in approximate relative position and different cells are denoted by color. Error estimation is standard deviation from the mean trace. **B.** Autocorrelelograms (ACGs) for example units from A. ACGs are normalized based on the sum of the ACG from 0 to 20 ms. **C.** Projection of waveforms on 2 principle components used for sorting. D) ACGs for each SU used for analysis displayed as heat map. ISI ratio (ISI_0-2ms_/ISI_0-20ms_) for all SU were < 0.1. **E.** Peristimulus time histogram response to short-mask (2 × 2 ms pulses) for all units recorded. Laser stimulation initiated at t = 0 ms. PSTHs are baseline subtracted, normalized to their maximum response bin value, and smoothed by convolving 1 ms binning with a Gaussian (σ = 4 ms).

**Supplementary Figure 3.**
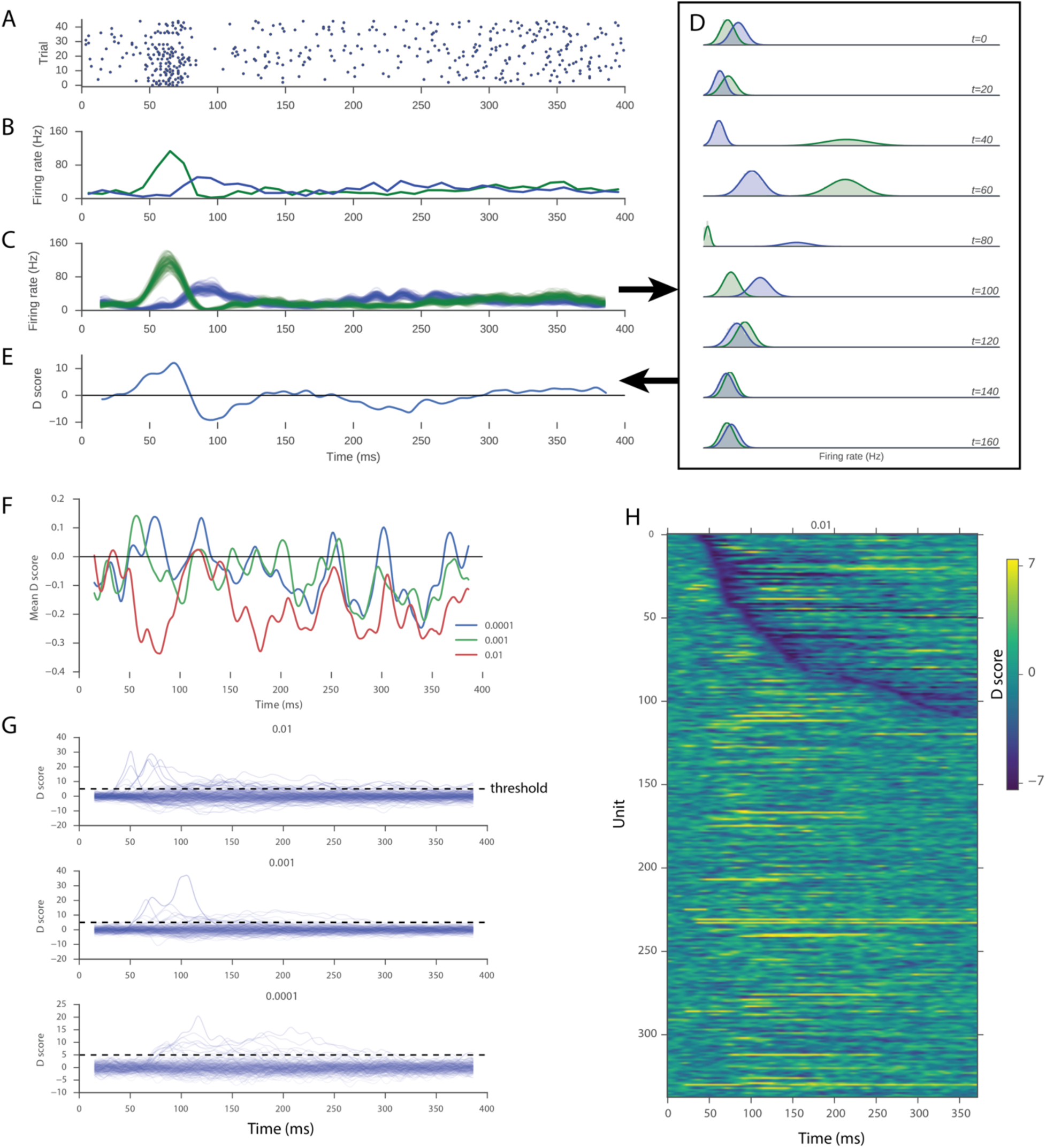
Defining odor response latencies across concentration. **A.** Raster of odor responses for example unit. Time 0 is defined as inhalation onset. **B.** PSTH plot from raster. Odor response PSTH is shown in green and blank response in blue. **C.** Bootstrapped PSTH plots. Each PSTH instance was created by sampling with replacement from trials, summing the responses across trials for each bin, and convolving a Gaussian kernel over this array (see methods). **D.** Histograms of bootstrapped responses for odor and blank trials at different time points relative to inhalation onset. Fits of normal distributions used to calculate D-scores are overlaid. **E.** D-score over time. Positive values specify excitation (odor response distribution is greater than blank distribution) and negative values specify inhibition relative to unit’s blank response. **F.** Mean D-score odor response across all units for each concentration tested. Note early inhibition at highest concentration. **G.** Individual unit D-score odor responses for all units. Each axis represents responses at the concentration specified above the axis. The threshold used to determine response latencies is overlaid as a dashed line. The threshold is constant across concentrations. **H.** D-scored responses of recorded population for one concentration of odorant. Units are sorted by latency to inhibitory response. Latency is determined by the first time point in which each units’ odor response D-score crossed a threshold of -5.

**Supplementary Figure 4.**
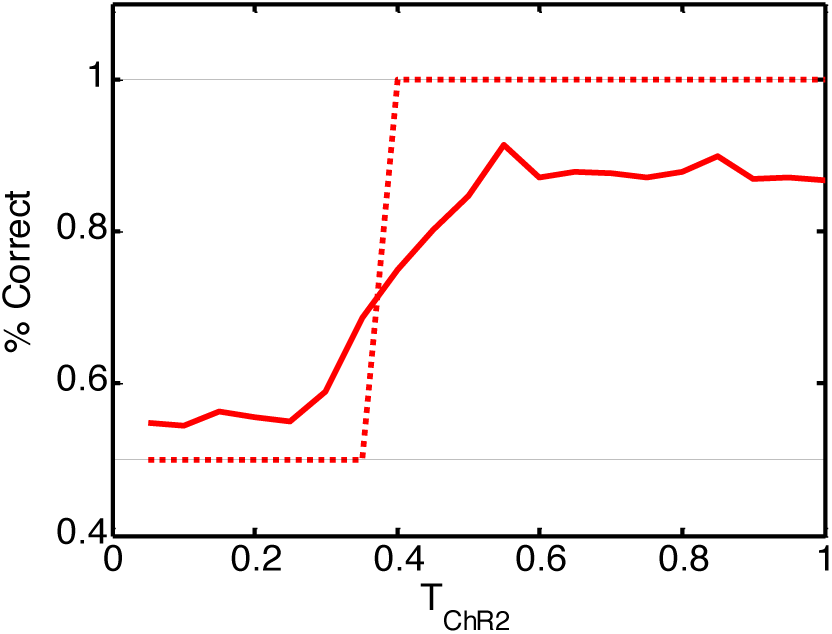
In our computational model, an incomplete suppression of animals’ behavior to 50% by the masking stimulus at short masking times is due to sources of variability and noise. Solid red line: same as in Figure 5D. Dotted figure, the same with no neural noise and for a single network weight configuration (emulating the conditions of a single animal). The latter modification was made to eliminate the variability due to varying performance in different animals. The behavior as affected by the mask jumps from pure 50% and 100% supporting the interpretation that, in the model, the deviations from pure 50% performance are due to internal sources of noise and individual variabilities.

**Supplementary Figure 5.**
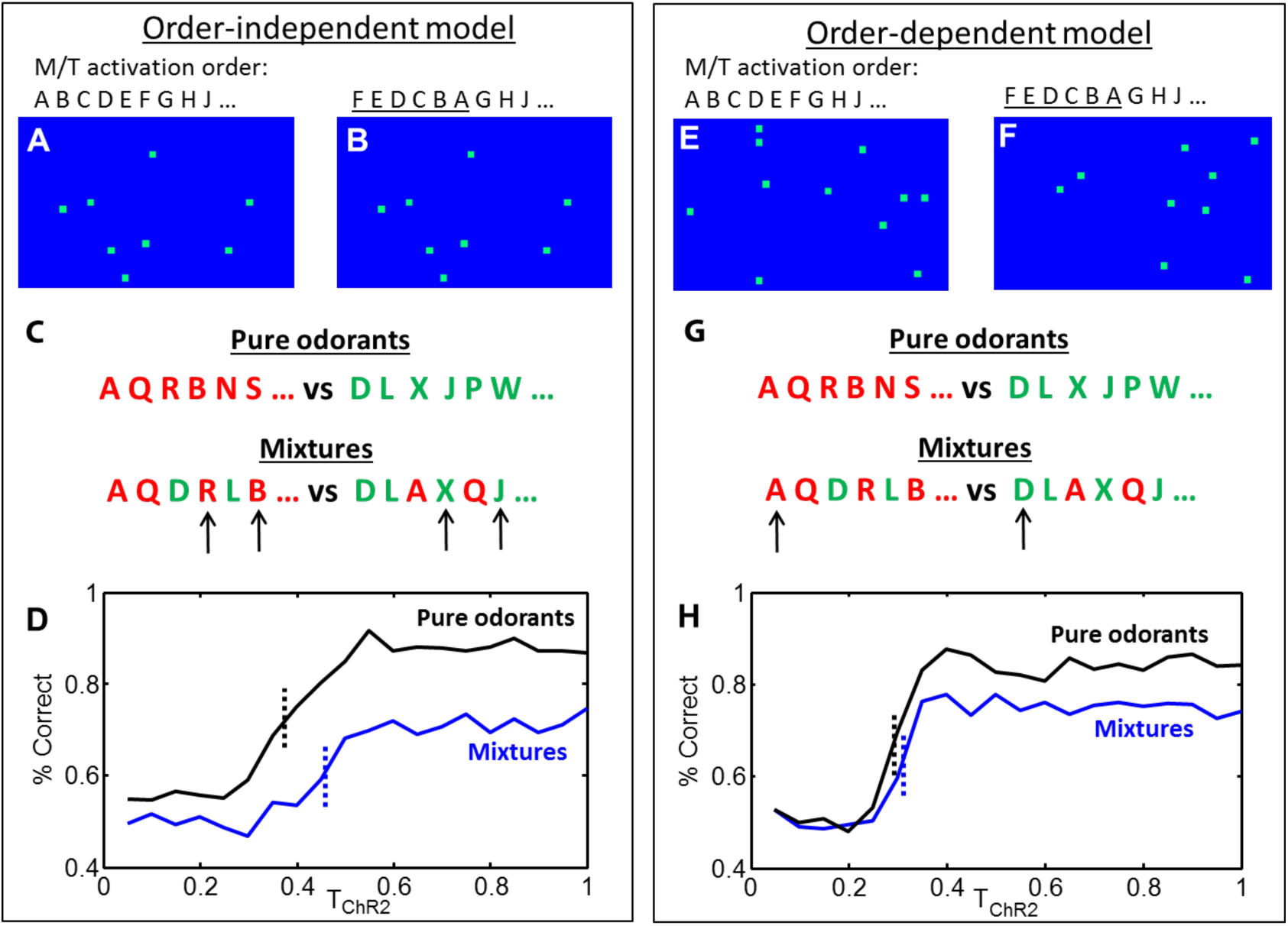
Order dependent and order independent primacy models give different predictions for the dynamics of masking response. **A-B.** Order-independent model. The patterns of PC activation are the same for different MT activation orders (letters on tops of panels), differing in the early stages of the sequence (differences underlined). These two glomerular activation sequences were chosen as examples to demonstrate the recruitment order independence of this instance of the model. **C.** Glomerular activation sequences in the modelling of masking response. To test the performance of the model in response to mask, we presented two different random sequences for pure odorants (top). For the case of mixtures, the recruitment order was obtained by mixing two pure odor sequences with the temporal offset equal to two positions. The sequence corresponding to the lower concentration component (40%) was delayed w.r.t. the higher concentration sequence (60%). Arrows show differences between pools of activated glomeruli. They occur later in the sequence (4^th^ and 6^th^ positions) delaying the effect of mask in the case of similar mixtures. **D.** Order independent model displays a small delay in the masking response for the case of mixtures (blue vs. black dotted lines). **E-F.** Order dependent model displays differing patterns of PC activation when the order of MT cell activation is different. To implement the sensitivity of the model to the order of activation sequence, we increased the strength of PB->PC weights from 0.15 to 0.3. This makes PC start detecting coincidences earlier in the sniff cycle. **G.** The activation sequences in this model were taken to be the same as in the order-independent model. Yet, the differences in the activation sequence emerge earlier in the sniff cycle (arrows) leading to earlier psychophysical performance. **H.** The order-dependent model shows a much smaller difference in the timing of behavioral responses to mask. The comparison of order-independent and order-dependent models suggests that the slight delay observed in the performance in similar mixtures of enantiomers (Fig. 3H) may be due to a form of order-independence in the decoding of the primacy sequence.

